# The Object Space Task shows cumulative memory expression in both mice and rats

**DOI:** 10.1101/198382

**Authors:** Lisa Genzel, Evelien Schut, Tim Schröder, Ronny Eichler, Angela Gomez, Irene Navarro Lobato, Francesco Battaglia

**Author notes:** equal contribution (Evelien Schut, Tim Schröder). corresponding author, Donders Institute, Radboud University and Radboudumc, Nijmegen/Netherlands.

## Abstract

Declarative memory encompasses representations of specific events as well as knowledge extracted by accumulation over multiple episodes. To investigate how these different sorts of memories are created, we developed a new behavioral task in rodents. The task consists of three distinct conditions (*stable, overlapping, random*). Rodents are exposed to multiple sample trials, in which they explore objects in specific spatial arrangements. In the *stable* condition, the locations are constant during all sample trials; in the test trial, one object’s location is changed. In the *random* condition, object locations are presented in the sample phase without a specific spatial pattern. In the *overlapping* condition, one location is shared (overlapping) between all trials while the other location changes during sample trials. We show that in the *overlapping* condition, instead of only remembering the last sample trial, rodents form a cumulative memory of the sample trials.

Here we could show that both mice and rats can accumulate information across multiple trials and express a long-term abstracted memory.

## INTRODUCTION

Memories are stored and retrieved in different ways, depending on the age of memory and the character of memorized information. In episodic memory, the details of the memorized event are retained. Conversely, semantic memory extracts general knowledge across multiple events. Memory consolidation processes may promote a transition between these two types of memory organization (1, 2). However, many tasks, especially for rodent subjects, cannot differentiate between the two, even though this differentiation is critical to further our understanding of memory mechanism (3).

Most memory paradigms used in rodents can be trained in a short time (1-2 sessions), enabling one to determine exact timings of memory interventions. But many such protocols require aversive reinforcers, such as electrical shocks in fear conditioning or avoidance learning (4, 5), or other strong motivators such as water or food reward. Such learning incentives strongly drive the neuromodulatory systems, a fact often not considered in studies using these tasks (6). In contrast, object recognition paradigms make use of a rodent’s natural tendency to explore more novel items, thus allowing for the investigation of memory processes without an intrinsic, difficult to control, side effect on motivation and emotion (7-9).

Another critical factor determining influencing memory acquisition is the frequency of events the animal experiences. In some memory tasks, the animal is exposed to repeated training trials. Spatial memory tasks such as the watermaze consist of multiple sample trials for the rodent to learn the location of a hidden platform most commonly trained across multiple days (10). The radial arm maze requires animals to repeatedly sample baited arms and their memory performance is assessed by the number of errors, namely the frequency of unbaited arm visits within a given trial; again, days to weeks of training are needed for the animal to perform above chance level (11, 12). Similarly, in some aversive conditioning paradigms, subjects undergo multiple pairings of a conditioned stimulus (CS) such as a tone or light with a mild foot shock (5). In other memory tasks, the animal only experiences a single event, which is the case in some fear memory paradigms, object recognition or object displacement memory (13–15). In object tasks, animals are allowed to explore two objects in a given environment for certain amount of time. After a delay, a short delay to assess short term memory or a delay of 24 hours to assess long-term memory, one of the objects is either replaced by a novel object (testing object recognition memory) or moved to a novel location (testing object position memory). Memory is assessed by calculating the difference in exploration time of the (for rodents preferred) novel item/location versus the familiar. In tasks where the number of events the animal experiences varies greatly, it is unclear which part of the training was significant to the animal’s performance. Is only the most recent event memorized by the animal? Or can memory be accumulated across extensive time periods or multiple trials?

These questions are key to understanding mechanisms of episodic vs. semantic memory (3), but they are difficult to address in most memory tasks. However, some recent work has attempted to study the accumulation of evidence across multiple events. An example is a modified version of the watermaze, in which evidence accumulation was assessed as mice were trained on multiple platform locations that were drawn stochastically from a specific spatial distribution and retrieval of ‘averaged’ memory of the learned platform locations was assessed after a 1-day or 30-day delay (16). Another example is paired-associate learning in rodents, in which memory of flavor-place associations is gradually learned with repeated trials over weeks, but later updating can occur within one trial (17–19).

Training procedures in these cumulative memory tasks are often lengthy and labor-intensive. In addition, information can be mainly acquired from either retrieval and/or updating: encoding and consolidation processes are difficult to study. A water-based paradigm such as that of Richards et al (16), is ill-suited for electrophysiological recording of brain activity during learning. We overcame these limitations by developing a task designed to extract information from multiple, similar events accompanied by suitable control conditions. The task is a variation on the traditional object-place memory task, exploiting a rodent’s natural tendency to explore novel configurations. In the *Object Space* task, we manipulate the stability of different components of the experience (here specifically: object position), making them more or less amenable to accumulation across episodes. We developed task versions that are suitable for both rats and mice.

In this new task, rodents are allowed to explore two objects, presented, across multiple trials in a *stable, overlapping* or *random* sequence. In the *stable* condition, objects are always presented in the same location across sample trials (see figure 2 and 3). In the test trial, after a delay, one object is moved to a novel location. We expect to see a preference for the object in the novel location in the test trial but no preference for either location over the course of training. This condition can be solved by remembering only the final sample trial (that is, using episodic memory or familiarity) or by creating a cumulative memory of all sample trials. The *overlapping* condition is our key condition. One object location remains stable across sample trials whereas the other object moves between one of three locations each sample trial. Importantly, the last sample trial shows the same configuration as the test trial after a delay. Thus, if the animal only remembers the most recent event it experienced, it will show no preference for either object location since both locations are familiar to the animal. Conversely, if the animal has accumulated the overlapping information of the stable location over time, it will still show a preference for the location less often shown. The control *random* condition consists of objects presented in random spatial configurations in which no patterns can be extracted and no place preference should develop.

The three conditions can be repeated multiple times in the same animals, thereby allowing for within-subject designs. Further, it is easy to combine behavioural training with physiological measures such as electrophysiology and other manipulations. The rat version of this task requires only one day of training involving 5 trials with 50min inter-trial intervals, a 10min test trial follows 24hrs later. In mice, this training protocol is repeated over the course of 4 days and a test trial follows 24hrs after the fourth training day. We further, ran a 4-week training paradigm in mice and could show that cumulative memory was retained for at least 5days. We show here that both rats and mice can express a cumulative memory.

## METHODS

### Subjects

Male C57Bl6/J mice, 7-8 weeks of age at the start of behavioral training (Charles River) and male Lister-Hooded rats (12 weeks, Charles River) were group housed with *ad libitum* access to food and water. Animals were maintained on a 12h light/dark cycle and tested during the light period. In compliance with Dutch law and Institutional regulations, all animal procedures were approved by the Central Commissie Dierproeven (CCD) and conducted in accordance with the Experiments on Animal Act.

### Behavioral training

#### Habituation

Animals were thoroughly handled in their second week after arrival in the animal facility. Each animal was actively handled daily for at least 5 minutes. We emphasize here that handling of the animals is extremely important. Picking them up by the tail is aversive and inadequate handling can affect the animal’s performance on multiple tasks (Gouveia and Hurst 2016). Mice and rats were handled so that they typically climbed by themselves on the experimenter’s hands when taking them out of the home cage and out of the training arena (see handling video https://www.memorydynamics.org/#/animal-handling/). After handling, animals were habituated to a square arena (75cmx75cm) for 5 sessions within 5 days. The walls and the floor were white or green to facilitate background subtraction in video analysis software. On the bottom side of the floor, magnets were placed in 4 locations for easy and consistent placement of the objects; objects were affixed to square metal plates (Fig 1). In the first habituation session, the animals were allowed to explore the box together with all cage mates for 30 minutes. In the second and third session, they were placed in the box individually for 10 minutes. In the final two sessions of habituation, two objects (towers made from Duplo blocks, not used in main experiment) were placed in the box at locations not used during training and the animals were allowed to explore for 10 minutes.

**Figure 1.**
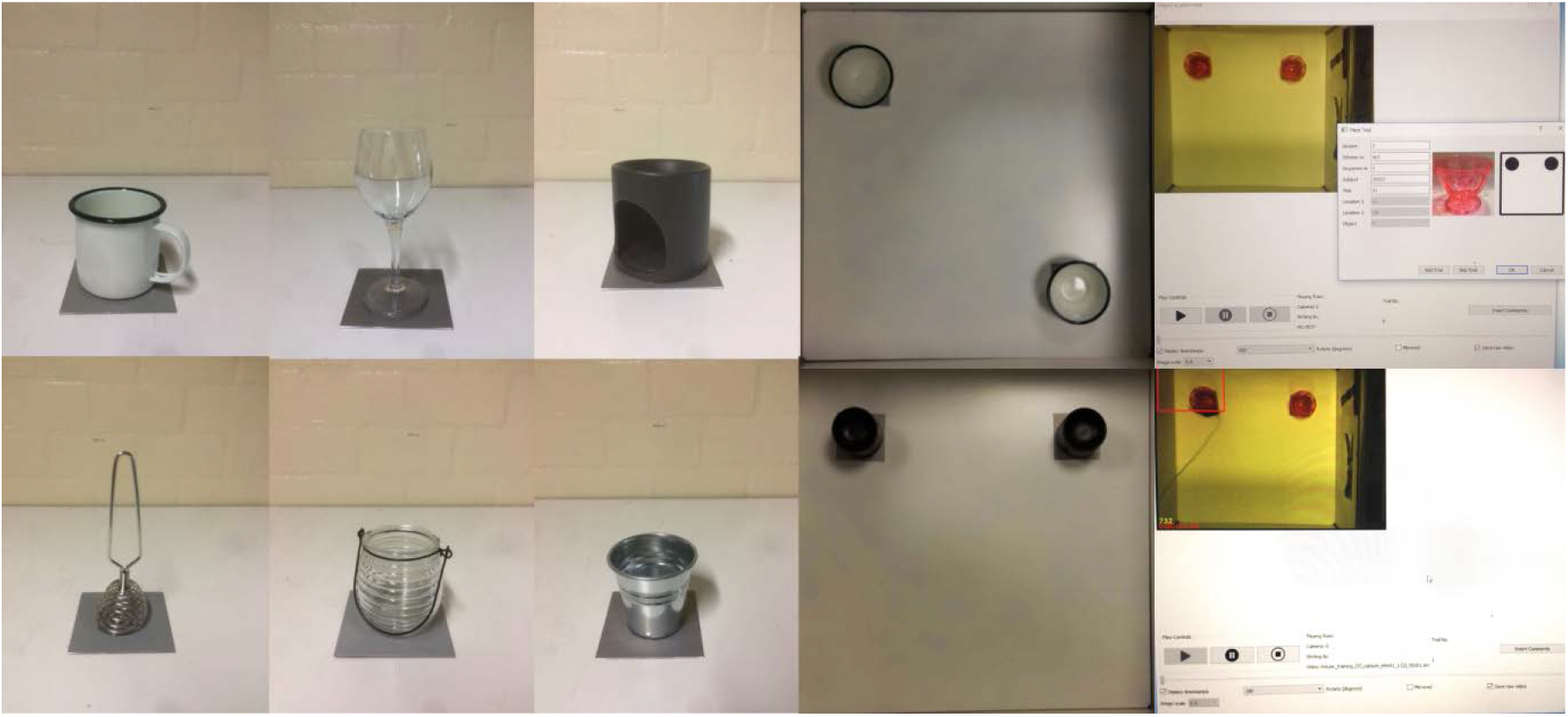
Object Space Task Materials: Examples of objects used in the object space task. Objects vary in size, width, texture and material. Objects were placed in two of the four corners. On the right: example of the object scorer program with pop-up pre-trial (top) and with scoring (bottom).

#### Training

The object space task consists of three conditions: stable, overlapping and random as described above (see Fig 2, 3). Conditions and locations were counterbalanced among animals and sessions and the experimenter was blinded to the condition. At the beginning of each session (2 days for rats, 5 days for mice), cues were placed on the walls inside the box, distributed intentionally non-symmetric. Thus, cues were typically not placed in the middle of each wall but would rather be distributed in a way that one cue would for example cover the lower left part of the wall while another cue would occupy the top right part of another wall. At least one 3D cue was placed above any of the walls to facilitate allocentric processing during the task. All cues were chosen to be high contrast and varied from session to session in general shape and geometry, to cater to the bad vision of rodents. A camera was placed above the box to record every trial and to allow for online scoring of exploration time. Multiple experimenters were involved in the experiment and each separate batch of animals (n=8 per batch) was trained by either one constant experimenter or by at least 2 experimenters in a rotational schedule, these difference had no effect on the replicability of the results from one batch of animals to the next.

**Fig 2.**
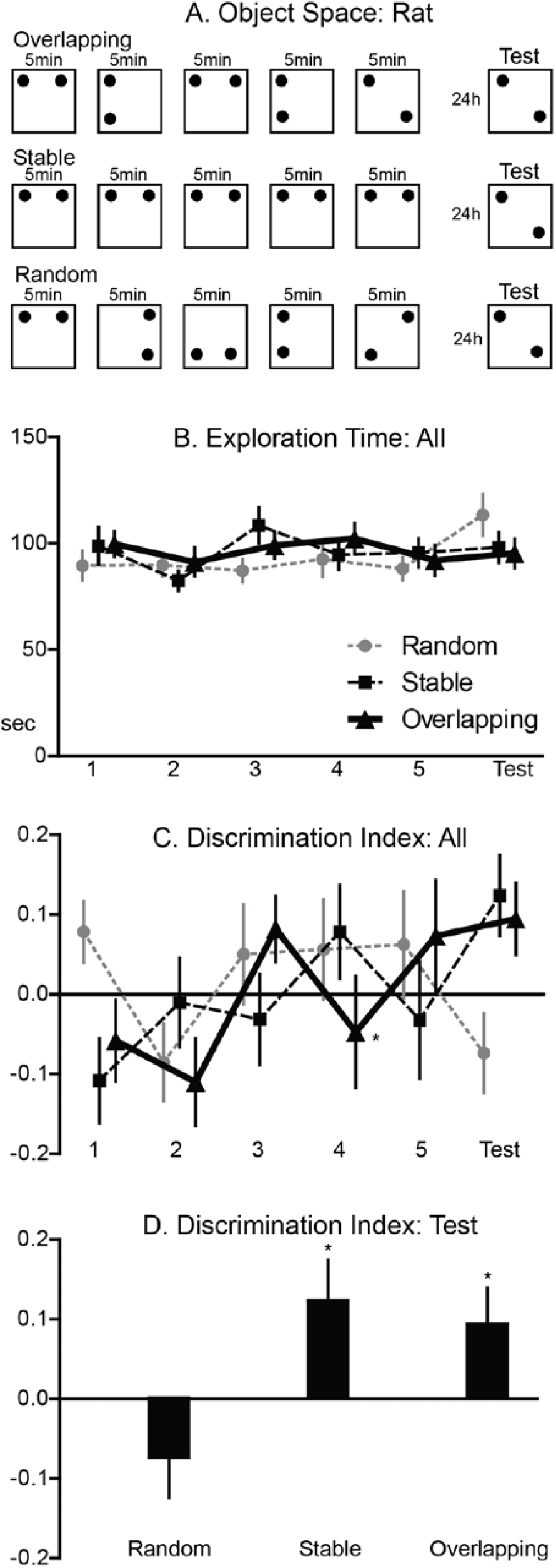
Object Space Task Rat: A. Panel: Trial structures for the three different conditions. In the *overlapping* condition, one location remains constant across all sample trials and the test trial, the second location varies. The locations in the last sample trial and in the test trial are equal, thus only cumulative memory across trials will lead to a preference for the less often shown location. In the *stable* condition, the locations remain the same in all sample trials and one object is displaced in the test trial. In the *random* condition, the locations were pseudo-randomly chosen to not allow extraction spatial patterns. One session consisted of 5 sample trials on one day and a test trial 24hrs later. B. Panel: Exploration time. The total exploration time remained constant across conditions and but a significant effect of trial was observed (p=0.017). C. And D. Panel: Discrimination Index (for statistical details see main text). The DI across sample and test trials showed a significant condition x trial interaction effect (p=0.011). In the test trial, there was a significant condition effect (p=0.008) and the DI was significantly above chance only for stable and overlapping condition (* p<0.05).

In each condition, animals were allowed to explore two objects for 5 minutes with an inter-trial interval of 30min for mice, 50min for rats. Mice were trained interleaved in groups of 4 with two groups per day (morning/afternoon), rats in groups of 8 (one group per day). Before the beginning of each sample trial, the box and the objects were thoroughly cleaned with 70% ethanol. Each sample trial consisted of a different pair of matching objects varying in height width, texture and material (including metal, glass, hard plastic and lacquered wood, see Fig 1 for example objects). Object sizes ranged from 4-26cm in height to 5-18cm in width. Objects were glued onto metal coasters and placed onto the magnets that were fixed on the floor of the arena.

Objects were never repeated during the training period of one condition (1 session). Rats received 5 sample trials in total. This procedure was repeated in mice over the course of 4 consecutive days in which they were presented with either 3 sample trials per day (see supplemental materials) or 5 sample trials per day, thus accumulating in 12 or 20 total sample trials. The test trial, 24hrs after the last sample trial, consisted of again two objects and animals were allowed to explore for 10 minutes, however only the initial 5 min were used (for 10 min results see supplemental materials).

In each species 4 batches of each 8 animals were run, resulting in a total of 32 animals. In mice one animal was excluded after running the first experiment (3-trial version) due to exploration times of less than 5sec and never experienced the 5-trial version. In rats, one animal was excluded due to exploration times of less than 5 seconds and the data of another animal was not included due to false placement of the objects during the test in the overlapping condition (not included in the other conditions due to the within-subject analysis).

Additionally, 8 mice were run on a 4 week version of the *overlapping* condition, with 3 weeks of each 25 trials (5/d, Mo-Fri) and a final trial on Wednesday of week 4. The second weeks Monday (trial 26) as well as the final trial (trial 76) were run with the same configuration as the previous sample trial to function as 3d and 5d test trials. All animals were included in the analysis.

#### Data acquisition

We developed an in-house program for training and scoring. The Object Scorer reads in previously prepared training sheets with the object and locations for each trial of each animal, presents this information at the beginning of each trial to the experimenter (see Fig 1) and automatically extracts exploration times from the manually-scored videos. Therefore, the operator cannot keep track of which animal is in which condition, and which is the stable vs. moved object for each trial, and he or she can be considered blind. Source code for the Object Scorer software is available at https://github.com/MemDynLab/Score.

#### Statistical analysis

The discrimination index (DI) used to assess memory performance was calculated as the difference in time exploring the novel object location and stable location divided by the total exploration time. This results in a score ranging from −1 (preference for the stable location) to +1 (preference for the moving object location). A score of 0 indicates no preference for either object location. Total exploration time and discrimination index were assessed with repeated measure ANOVAs with factors condition and trial (6 trials) in rats. Additionally, in rats the DI of the test trial was assessed with a repeated measure ANOVA for condition. Due to the different training schedule in mice (5 trials per day for four days but only one test trial on day 5) the sample trials were separately assessed by including the factors condition, trial, and day across the 20 sample trials in mice in an repeated measure ANOVA. To test long-term memory in mice the final sample and test trial were included in a repeated measure ANOVA with factors condition and trial. When a significant main effect or interaction was found, one sample t-tests were performed to analyze memory performance with respect to chance level in the last sample trial and test trial.

The 4-week *overlapping* training was analyzed with repeated measure ANOVA for week (each animal averaged across the week) for exploration time and discrimiation index. Further, the two test trials (3d and 5d) were tested to chance level with a one-sample t-test.

## RESULTS

### Rat training: 2-day training paradigm

Rats were trained for 5 trials in one day before being retested 24h later (n= 30, Fig 2). There was a small but significant trial effect on total exploration time but no condition effect or conditionXtrial interaction (condition F_2,58_=0.27, p=0.76; trial F_5,145_= 2.87, p=0.017; conditionXtrial F_10,290_=1.38, p=0.22). The discrimination index (DI) across all 6 trials showed a significant conditionXtrial interaction but no significant condition effect and only a marginal significant trial effect (condition F_2,58_=0.04, p=0.96; trial F_5,145_=2.19, p=0.059; conditionXtrial F_10,290_= 2.35, p=0.011). When focusing on the final test trial, there was a significant condition effect (condition F_2,58_=5.29, p=0.008). In addition, memory performance at the stable condition was significantly above chance (stable: t_29_=2.44, p=0.021). In addition, DI for the overlapping condition was also significantly above chance (*overlapping*: t_29_=2.09, p=0.045). This was in contrast to random (t_29_=-1.47, p=0.15). Thus, we can show that both overlapping and stable training conditions led to significant memory expression at test 24h later with preferred exploration of the less stable object location, which was not seen in the random condition.

### Mouse training: 5-day training paradigm

Mice were trained with 5-trials a day across 4 days, with a test 24h later (n=31, Fig 3). A 3-trial version was also piloted (n=7, see supplemental materials). No differences in total exploration time were found between conditions or any interaction with condition but significant trial and day effects were seen during the 20 sample trials (condition F_2,60_=0.51, p=0.59; trial F_4,120_=15.25, p<0.001; day F_3,90_ =9.98, p<0.001; conditionXtrial F_8,240_=0.42, p=0.85, conditionXday F_6,180_=0.35, p=0.85; Fig 3). In addition, there was a significant trialXday interaction on exploration time (trialXday F_12,360_=7.09, p<0.001) but importantly no 3-way interaction (conditionxdayxtrial F_24,720_=0.75, p=0.68). Discrimination Index across sample trials (20 trials) showed a marginal significant trial effect and more importantly a significant trial x condition interaction (condition F_2,60_=0.52, p=0.52; trial F_4,120_=2.0, p=0.093; conditionXtrial F_8,240_=2.3, p=0.042), indicating that only in the overlapping condition a build-up was seen during the 5 trials each day. All other main and interaction effects were not significant (day F_3,90_=0.93, p=0.43; conditionXday F_6,180_=0.87, p=0.52; conditionXdayXtrial F_24,720_=1.08, p=0.38), except for a significant trialXday interaction (F_12,360_= 1.97, p=0.026). Concerning the final sample and test trial, there was a significant trialXcondition interaction effect (condition F_2,60_=2.0, p=0.14, trial F_1,30_=0.12, p=0.73; conditionXtrial F_2,60_=0.16, p=0.046). One sample t-tests indicated memory performance above chance for the stable condition at test (t_30_=3.0, p=0.005). Further, memory performance on both the last sample trial and test in the overlapping condition was significantly above chance, indicating that mice accumulated memory over the course of training, which led to long-term memory expression at 24h (final sample trial t_30_=3.0, p=0.005; test t_30_=2.16, p=0.039). Finally, no significant effects were observed in the random condition (final sample trial: t_30_=0.68, p=0.5; test trial: t_30_=-0.34, p=0.73). Thus, as in rats, both overlapping and stable training conditions led to significant memory expression at test 24h later with preferred exploration of the less stable object location, which was not seen in the random condition.

**Fig 3.**
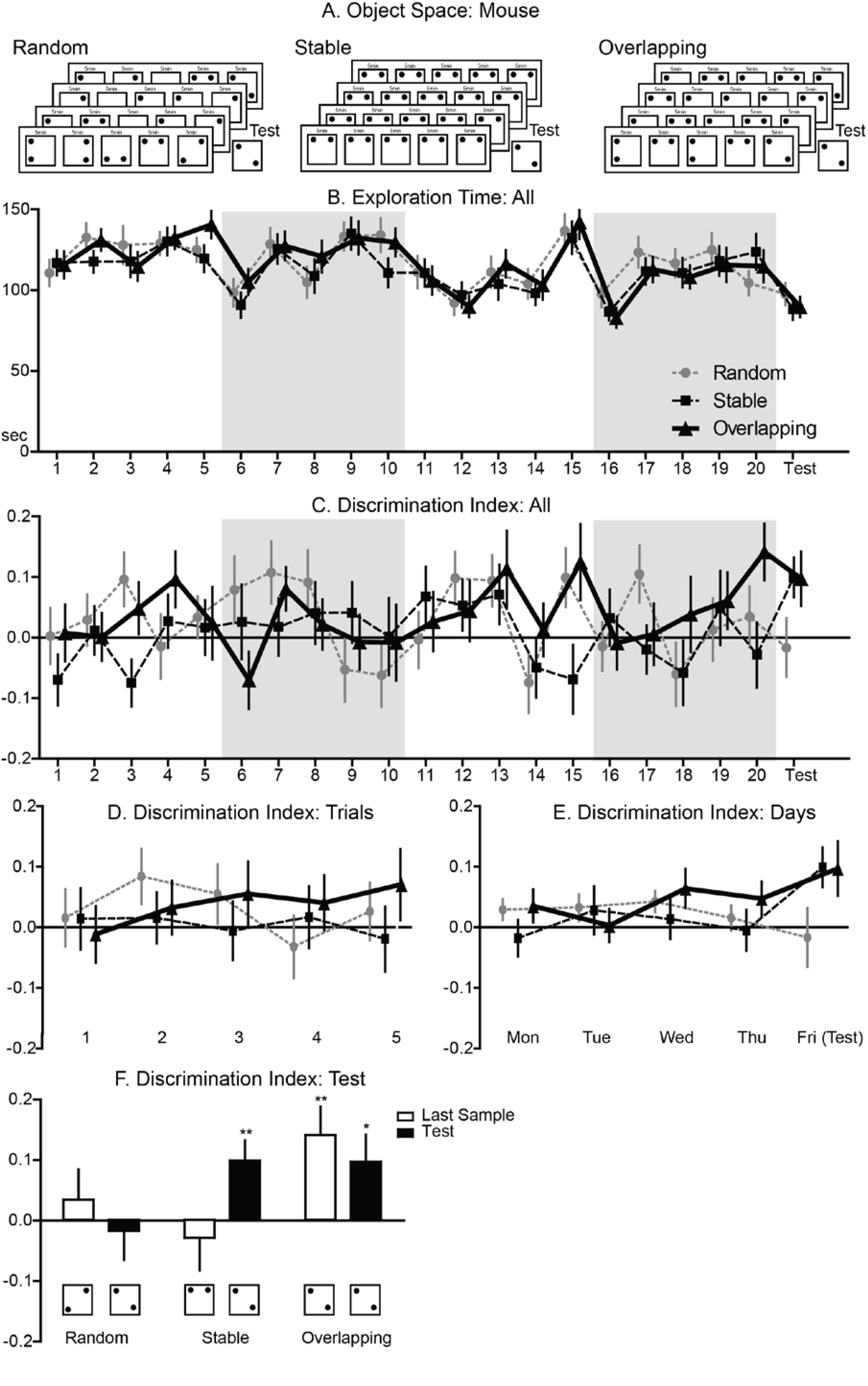
Object Space Task Mouse: A.Panel: Trial structures for the three different conditions. In the *overlapping* condition, one location remains constant across all sample trials and the test trial, the second location varies. The locations in the last sample trial and in the test trial are equal. In the *stable* condition, the locations remain the same in all sample trials and one object is displaced in the test trial. In the *random* condition, the locations were pseudo-randomly chosen to not allow extraction spatial patterns. One session consisted of 5 sample trials for 4 subsequent days and test trial 24hrs later. B. Panel: Exploration time over the course of all 20 sample trials and test trial for each condition. Alternating white and grey shaded areas indicate the individual training days and test day. The total exploration time per sample trial remained constant across conditions, however significant effects of trial, day and a significant trialXday interaction were observed (condition p=0.59; trial p<0.001; day p<0.001; trialXday p<0.001). C. Panel: Discrimination Index for all 20 sample trials and test trial across conditions. Alternating white and grey shaded areas indicate individual training days and the test day. D. Panel: Discrimination Index per sample trial over the course of all four training days across conditions. A marginal significant effect for trial has been found (p=0.09). More importantly, a significant conditionXtrial interaction was observed (p=0.042), indicating only a build-up of preference for the less stable location over the daily trials in the overlapping but not stable or random condition. E. Panel: Discrimination Index for each training day (the 5 sample trials for each training day averaged) and test day per condition. F. Panel: Discrimination Index at the final training trial and test trial, which showed a significant trialXcondition interaction (p=0.046). Memory performance was significantly above chance level in the overlapping condition for both the last sample trial and test trial (last sample p<0.01; test p<0.05). In the stable condition, only the test trial showed a significant effect (last sample p=0.59; test p<0.01). No significant effects were observed in the random condition (last sample p=0.50; test p=0.73).

### Mouse training: 4-week training paradigm

To test if our *overlapping* training led to a memory representation that lasts longer than 24h, eight mice were additionally trained with 5-trials a day across 25 days, with a final test 5 days later (n=8, Fig 4). Both the second week’s Monday (trial 26) as well as the final trial (trial 76) were run with the same configuration as the previous sample trial to function as 3d and 5d test trials. Exploration time remained stable (week F_2,14_=0.4, p=0.96) and the discrimination index remained positive indicating decreased preference for the stable location in our *overlapping* condition (week F_1.2,14_=0.5, p=0.53). The 3d and 5d test showed that even after longer periods the abstracted memory representation is still expressed with both tests above chance level (3d t_7_=2.7, p=0.033; 5d t_7_=3.7, p=0.008).

**Fig 4.**
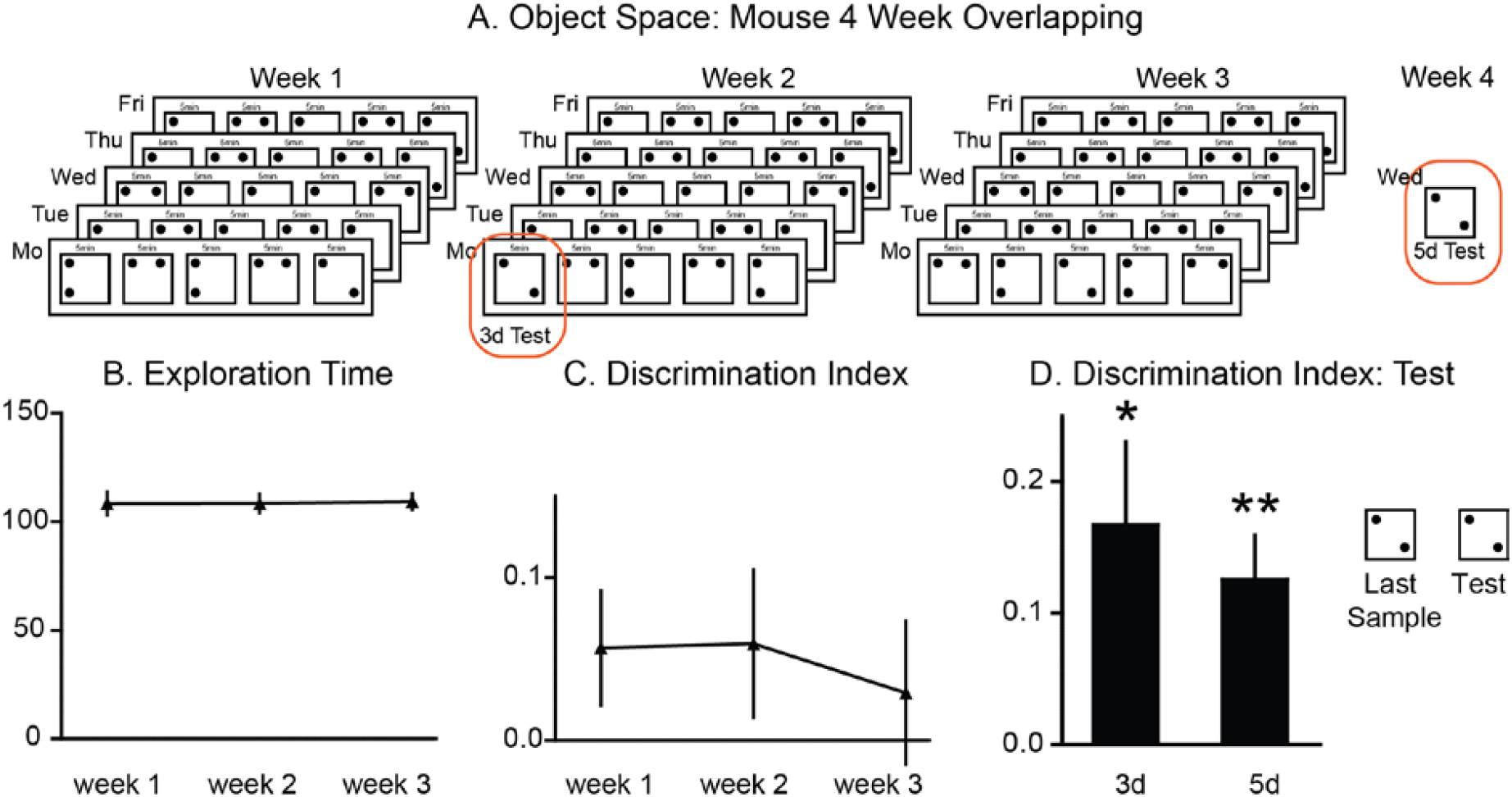
Object Space Task Mouse – 4 week overlapping training: A Panel: Trial structures for the four week version of the *overlapping* condition. Across the four weeks, one location remains constant across all sample trials and the test trial, the second location varies (5 trials per day, 5 days for the first three weeks). The first trial on Monday in week 2 (trial 26) as well as the final trial on Wednesday in week 4 (trial 76) function as 3d and 5d test trial respectively. B Panel: Exploration Time remained stable across the three weeks (p=0.96). C Panel: The discrimination index remains stable with preference for the less often shown location across the three weeks (p=0.5). D Panel: Test To control for episodic memory effects the locations in the last sample trials and in the tests trial are equal. Both 3d and 5d after training the animals show a significant cumulative memory effect with preference for the less often shown location (*p=0.033, **p=0.008).

## DISCUSSION

Knowledge extraction is a gradual process that requires the experience of multiple, similar (or overlapping) events, in contrast episodic memory is by definition based on one event (2, 20, 21). Although some tasks have previously been developed to study cumulative memory, we attempted to develop a task that is simple and easy to implement, that allows for a time-saving within-subjects design and makes use of a rodent’s natural behavior without any external motivators. We have successfully demonstrated that the *Object Space task* can be used to test for cumulative memory and contains both a positive control condition (the *stable* condition) that can be solved with a single event or recency memory as well as a random condition as negative control. By the end of training, both rats and mice show cumulative memory in the *overlapping* condition, indicated by a positive DI in the test trial. DI in the *stable* condition is, as expected, only biased in the test trial. Finally, object locations in the *random* condition were treated without preference by the animals.

Because the same configuration is used as in the last training trial, the test trial provides a control for any recent memory-like effects in our *overlapping* condition, clarifying whether the animal uses accumulated memory over the course of learning instead of their most recent experience to guide their behavior. If the animal shows no preference for either object location at the test trial, it can mean two things. Either the animal behavior is guided by remembering its most recent experience or the animal has not acquired a long-term, cumulative memory. Even though we cannot assume that the encoding strengths for the *stable* and *overlapping* condition are exactly the same, the *stable* condition does help to differentiate these two effects, since if the animal can retain a memory of the most recent experience but not a cumulative memory it still will be above chance in this condition. One could argue that the different training conditions result in different types of memories (e.g. abstracted for overlapping and episodic for stable) and thus a direct comparison with an ANOVA is not warranted and only a t-test to chance for each condition is critical to test for significant memory expression. Here, however, this distinction is of no importance since both approaches show significant results. Thus, all three conditions together *(overlapping, stable* and *random)* enables us to test if an animal under current conditions can remember an event and/or a cumulative memory. We further have expanded the approach in mice and showed, that the abstracted memory in the *overlapping* condition is not only expressed 24h later, but is retained for longer time periods as seen in the 3day and 5day tests.

While mice require multiple sample trials across multiple days to acquire cumulative memory in this task, rats require just one day of training consisting of 5 sample trials in total. Despite this difference in training duration and the definite slower learning curve in mice, we see it as an advantage that this task can be used in both rodent types. Several studies have compared performance in various (complex) tasks in rats and mice and often concluded that mice cannot perform as well as rats (22–24). However, as we show in our task, by adapting the protocol mice are able learn this task and retain the information over longer time periods, thereby expanding the opportunities for the use of this task in numerous animal models, taking advantage of the extensive molecular and genetic toolbox currently available for mice. Despite these differences in training duration, we expect that learning in this task underlies similar mechanisms in both rats and mice. However, we cannot draw any conclusions on this until further research has been conducted.

In addition to adapting the task to rats and mice, we developed a software to track the exploration behavior and allow for online scoring of exploration periods. The program automatically reads in pre-defined trial structures and only informs the experimenter about the objects and locations used right before each trial. Combining this approach with interleaved testing of several animals during one experimental session, we effectively blind the experimenter with respect to the condition in the current trial and therefore enable them to score exploration behavior online without introducing an experimenter bias.

In the future, this task will allow for the investigation of the neural circuits contributing to cumulative and event memory. In contrast to water-based paradigm such as that of Richards et al (16), this task is well-suited for electrophysiological recording of brain activity during learning (see Fig 5). Further, this task is especially suitable as memory conditions tapping into memory accumulation vs. event memory can be presented in the same spatial layout and with very similar overall behavior, as indicated by the lack of difference in total exploration time across conditions.

**Fig 5.**
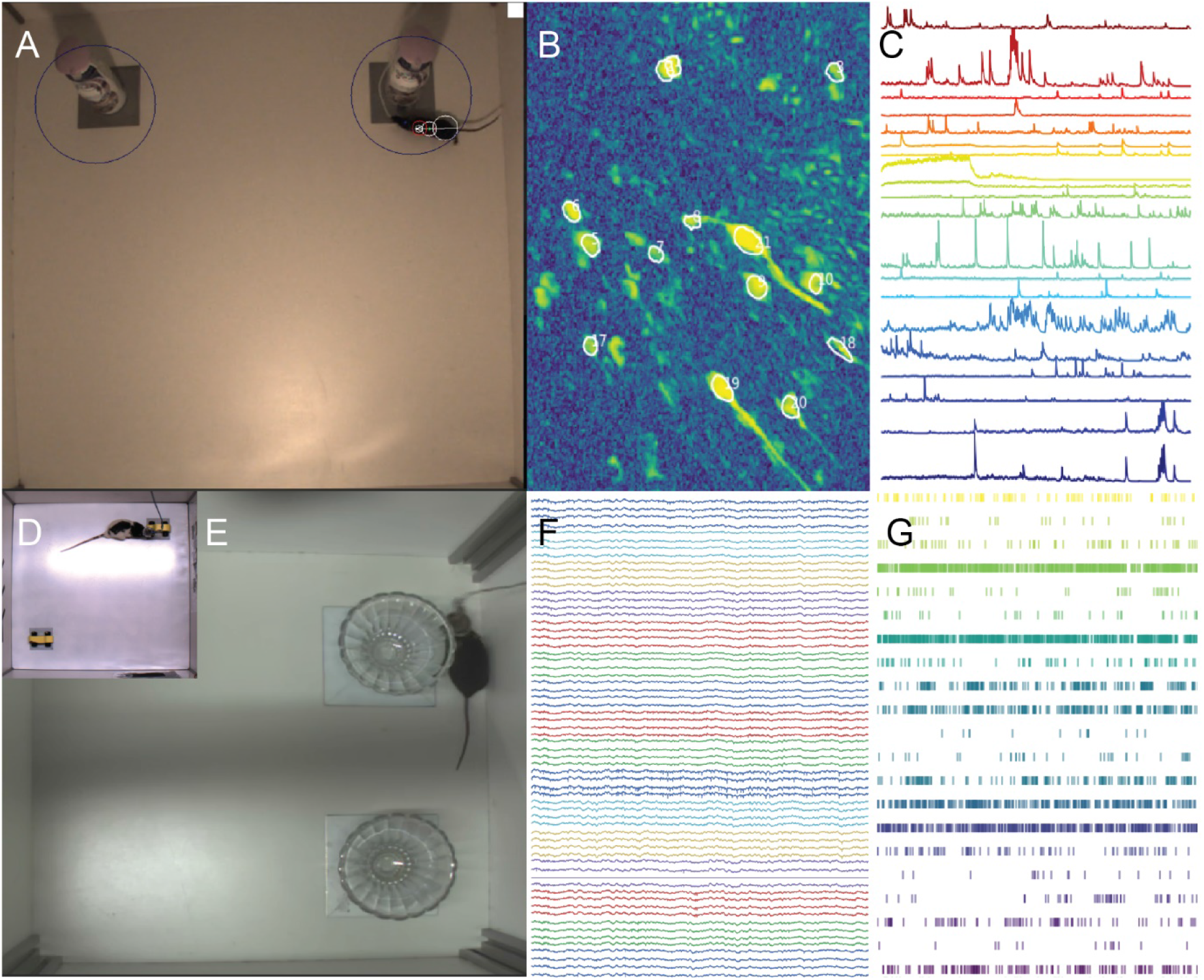
Object Space Task–Calcium Imaging and Electrophysiology: Example of a mouse running the OS Task with calcium imaging (Inscopix, A), example raw signal (B) and extracted calcium transients (C). Example of a rat (D) and mouse (E) running the OS task with electrophysiological implants, example raw LFP signal (F) and extracted unit activity (G).

Previous studies have provided evidence that the hippocampus is more involved in the processing of recent experiences that include episodic details, whereas the prefrontal cortex accumulates information from multiple, similar experiences, thereby creating a more stable but also more generalized memory over time (1, 17, 19, 25). We can hypothesize that successful performance on the overlapping condition involves the integration of multiple or all events in the prefrontal cortex, thereby creating a stable representation of the overlapping object location in space. While the classic version of our stable condition, namely 24h-object-displacement memory, is usually described as a hippocampal-dependent task (26–29), we cannot assume that our stable condition is also dependent on the hippocampus due to the increased number of sample phases. Object-displacement memory requires the animal to experience only one event, in the *Object Space task* the animal experiences multiple events of the same spatial configuration. Thus, the animal can solve this task by using both its most recent experience and the cumulative memory of the events.

In conclusion, the object space task can be used to study cumulative memory in both rats and mice. Rats require one day of training to acquire a cumulative memory while mice require multiple days of training in order to learn this task. Although we can speculate about a critical role of both prefrontal cortex and hippocampus to acquire cumulative memory in the object space task for both rodent types, the neural mechanisms underlying memory performance should be determined next.

## Acknowledgements

We would like to thank all students who have performed the Object Space task for this and other projects and who have convinced us of the reliability of the task: Gülberk Bayraktar, Nikkie Cornelisse, Hussein Ghareh, Federico Giuliani, Sidney Hulzebos, Joanne Igoli, Stefanos Loizou, Luc Nijssen, Luca Reinink, Olaf Stoutjesdijk, Minou Verhaeg, Thijs Groenveld, Thom Joosten, Liz van den Brand, Violeta Caragea, Robin Lempens, Rian Kraan, Koen van den Berg, Wessel Fanselow, Daan van den Tillaart, Nikki van den Berg, Milan Bogers.

## Supplemental Materials

### Mouse training: 3 sample trials per day

#### Results

Initially, we trained mice (n=7) with 3x 5min sample trials per day for each condition (Suppl. Fig 1). Exploration time of animals over the course of training was not significantly different between conditions but did show a significant trial effect (condition F_2,12_=0.8, p=0.47; trial F_12,72_=14.3, p<0.001; conditionXtrial F_24,144_= 0.56, p=0.95). Discrimination index averaged for each day showed a significant effect of condition (condition F_2,12_=6.2, p=0.014, day F_4,24_=0.84, p=0.51; conditionXday F_8,48_=0.6, p=0.78). Focusing on the final training and test trial, again a significant effect of condition was found (condition F_2,12_=4.6, p=0.033, trial F_1,6_=1.75, p=0.23; conditionXtrial F_2,12_=1.3, p=0.31). Performance in the *overlapping* condition was above chance at the final sample trial but not at the test trial (final training trial: t_6_=4.2, p=0.006; test trial: t_6_=1.2, p=0.27), Furthermore since we did not observe even a numerical effect on the *stable* condition (final training trial: t_6_=0.6, p=0.57; test trial: t_6_=-1.5, p=0.18, *random* final training trial: t_6_=-1.6, p=0.16; test trial: t_6_=-0.1, p=0.99), indicating that there was no 24hr long-term memory effect after training. Together these results suggest that more extensive training is needed for mouse subjects, therefore we chose to train the mice on 5 sample trials per day instead of 3.

**Supplemental Fig 1.**
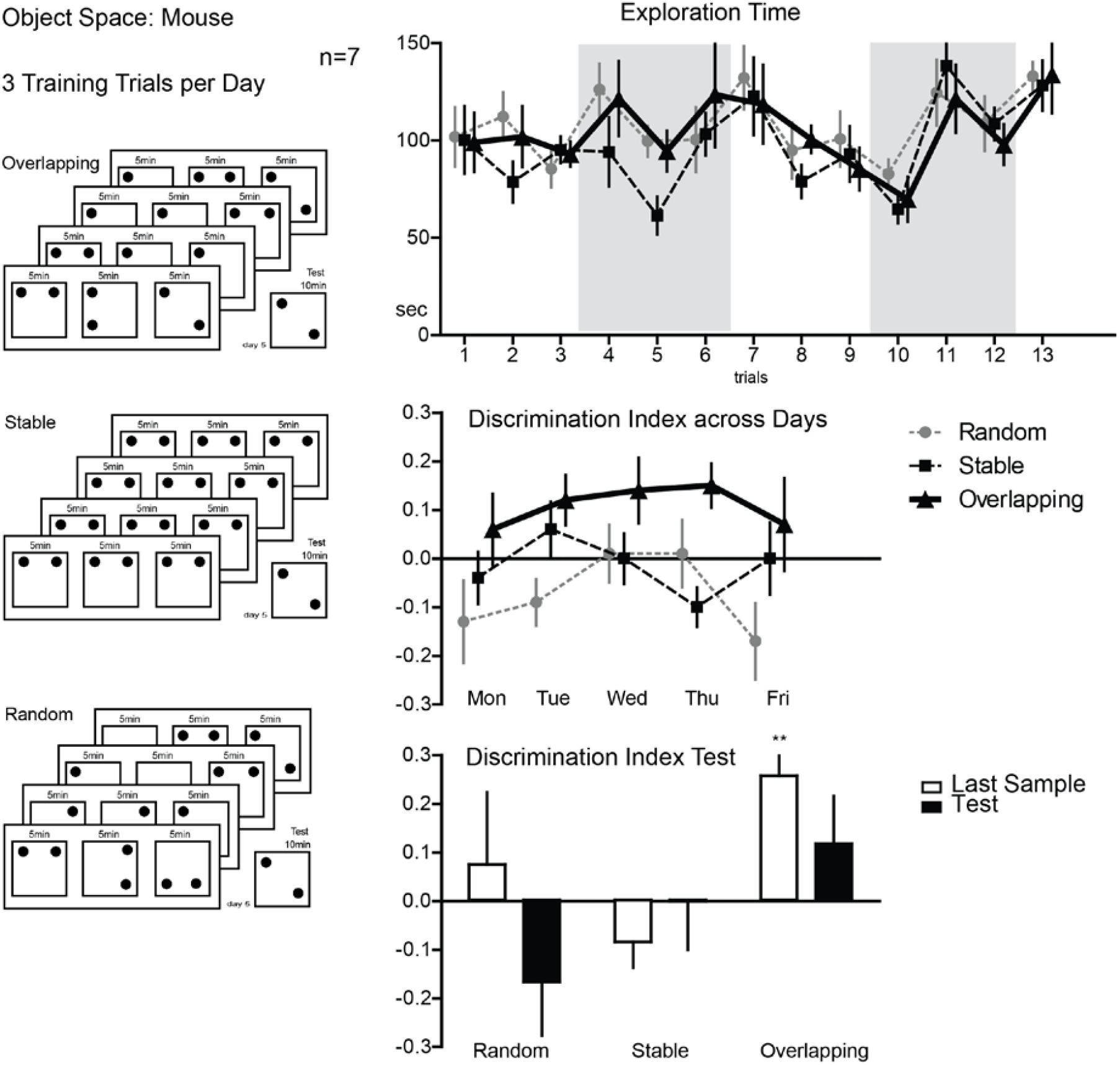
Object Space Mouse 3 Sample Trials Per Day: Left panel: Trial structures for the three different conditions. In the *overlapping* condition, one location remains constant across all sample trials and the test trial, the second location varies. The locations in the last sample trial and in the test trial are equal. In the *stable* condition, the locations remain the same in all sample trials and one object is displaced in the test trial. In the *random* condition, the locations were pseudo-randomly chosen to not allow extraction spatial patterns. One session consisted of 3 sample trials for 4 subsequent days and test trial 24hrs later. Top right panel: Exploration time over the course of all 12 sample trials and test trial per condition. Exploration time was not significantly different between conditions but did show a significant trial effect (trial p<0.001). Mid right panel: Discrimination Index averaged across training days and test day for each condition. Discrimination index averaged for each day showed a significant effect of condition (condition p=0.014). Bottom right panel: Discrimination Index at the final sample trial and test for each condition. A significant effect of condition was found (condition p=0.033). Performance in the overlapping condition was above chance at the final sample trial but not at the test (final sample trial: p=0.006; test: p=0.27), Furthermore, we did not observe a significant increase in memory performance on the stable condition test (final sample trial: p=0.57; test: p=0.18, random final sample trial: (p=0.16; test: p=0.99), indicating that there was no 24hr long-term memory effect after training.

### Rat and mouse training: 10min test

Animals were allowed to explore for 10 minutes during the test (Fig 2). Focusing on the test trial in rats, a marginal significant effect was found for condition (F_2,58_=2.64, p=0.08). Additional analyses showed no effects on the random condition (t_29_=-0.74, p=47). Further, the discrimination index for the stable condition was significantly above chance, indicating a 24hr memory effect, whereas no significant effects were found in the overlapping condition (*stable* t_29_=2.53, p=0.017, *overlapping* t_29_=1.58, p=0.13). However, in the 5min test we did find a significant effect in the overlapping condition (p<0.05). This indicates that rats spend more time exploring the moving object versus the stable object in the first 5 minutes of test, which clearly indicates a memory effect. However, in the last 5 minutes of the test, they have the tendency to return to exploring the stable object location more.

In mice, focusing on the last sample trial and test, a marginal trial X condition interaction effect was found (trail F_1,30_=0.06, p=0.8; condition F_2,60_=1.56 p=0.22; trial X condition F_2,60_=2.88, p=0.064). In addition, memory performance was significantly increased in the stable condition at test, indicating 24hr memory can still be observed with the 10min test (*stable* final sample trial t_30_=0.289 p=0.78; test t_30_=3.01, p<0.01; *random* final sample trial t_30_=0.68, p=0.5; test t_30_=-0.53, p=0.59). Performance at the final sample trial was significantly increased in the overlapping condition (t_30_=2.62, p=0.014) and a marginal effect was observed at test (t_30_=1.83,p=0.077). Thus, even though we can still observe the memory effects with a 10min test compared to a 5in test, the observed effects are stronger for the first 5 minutes compared to the whole 10min test.

Compared to the data from the 5min test, these results from both rats and mice indicate that analyses from the full 10 minutes of test are not able to demonstrate the memory effects as strongly as we observe during the first 5 minutes of the test. Both rats and mice spend more time exploring the object location that is more novel to them in the beginning of the test. Then the time spent exploring the familiar object increases as the 10min test progresses. Hence, a 5min test is a better representation of memory performance in the object space task.

**Supplemental Fig 2.**
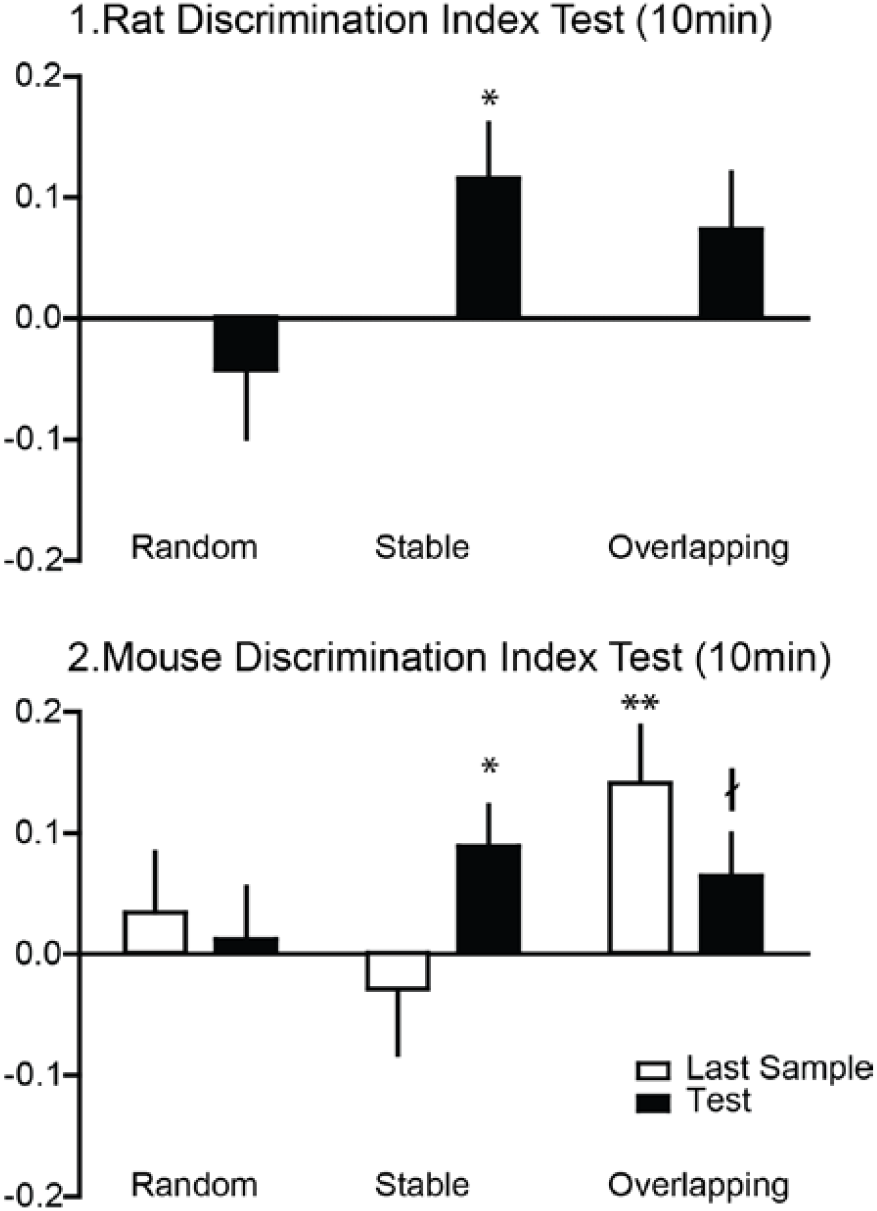
Discrimination Index At 10min Test: A. Panel: Rat Discrimination Index at 10 min test. Discrimination Index was calculated from 10 minutes of exploration during the test. A marginal significant effect was found for condition (p=0.08) with a significant increase in memory performance in the stable condition (p=0.017). B. Panel: Mouse Discrimination Index at lOmin test. Focusing on the final sample trial and test, a marginal trial X condition effect was found (p=0.064). In the stable condition, memory performance was significantly increased during test (p<0.01). Performance at the final sample trial was significantly increased in the overlapping condition (p=0.014) and a marginal effect was observed at test (p=0.077).

## REFERENCES

1. Frankland PW, Bontempi B. The organization of recent and remote memories. Nat Rev Neurosci. 2005;6(2):119–30. Epub 2005/02/03.

2. Moscovitch M, Cabeza R, Winocur G, Nadel L. Episodic Memory and Beyond: The Hippocampus and Neocortex in Transformation. Annu Rev Psychol. 2016;67:105–34.

3. Hardt O, Nadel L. Systems consolidation revisited, but not revised: The promise and limits of optogenetics in the study of memory. Neurosci Lett. 2017;5(17):30971–0.

4. Maren S. Neurobiology of Pavlovian fear conditioning. Annu Rev Neurosci. 2001;24:897–931. Epub 2001/08/25.

5. Tovote P, Fadok JP, Luthi A. Neuronal circuits for fear and anxiety. Nat Rev Neurosci. 2015;16(6):317–31. Epub 2015/05/21.

6. Martin-Soelch C, Linthicum J, Ernst M. Appetitive conditioning: neural bases and implications for psychopathology. Neurosci Biobehav Rev. 2007;31(3):426–40. Epub 2007/01/11.

7. Ennaceur A, Delacour J. A new one-trial test for neurobiological studies of memory in rats. 1: Behavioral data. Behav Brain Res. 1988;31(1):47–59. Epub 1988/11/01.

8. Ennaceur A, Neave N, Aggleton JP. Spontaneous object recognition and object location memory in rats: the effects of lesions in the cingulate cortices, the medial prefrontal cortex, the cingulum bundle and the fornix. Exp Brain Res. 1997;113(3):509–19. Epub 1997/03/01.

9. Warburton EC, Brown MW. Neural circuitry for rat recognition memory. Behav Brain Res. 2015;285:131–9. Epub 2014/10/16.

10. Morris R. Developments of a water-maze procedure for studying spatial learning in the rat. J Neurosci Methods. 1984;11(1):47–60. Epub 1984/05/01.

11. Olton DS. The radial arm maze as a tool in behavioral pharmacology. Physiol Behav. 1987;40(6):793–7. Epub 1987/01/01.

12. Bontempi B, Laurent-Demir C, Destrade C, Jaffard R. Time-dependent reorganization of brain circuitry underlying long-term memory storage. Nature. 1999;400(6745):671–5. Epub 1999/08/24.

13. Okuda S, Roozendaal B, McGaugh JL. Glucocorticoid effects on object recognition memory require training-associated emotional arousal. Proc Natl Acad Sci U S A. 2004;101(3):853–8. Epub 2004/01/09.

14. Bermudez-Rattoni F, Okuda S, Roozendaal B, McGaugh JL. Insular cortex is involved in consolidation of object recognition memory. Learn Mem. 2005;12(5):447–9. Epub 2005/09/17.

15. Beldjoud H, Barsegyan A, Roozendaal B. Noradrenergic activation of the basolateral amygdala enhances object recognition memory and induces chromatin remodeling in the insular cortex. Front Behav Neurosci. 2015;9:108. Epub 2015/05/15.

16. Richards BA, Xia F, Santoro A, Husse J, Woodin MA, Josselyn SA, et al. Patterns across multiple memories are identified over time. Nat Neurosci. 2014;17(7):981–6. Epub 2014/06/02.

17. Tse D, Takeuchi T, Kakeyama M, Kajii Y, Okuno H, Tohyama C, et al. Schema-dependent gene activation and memory encoding in neocortex. Science. 2011;333(6044):891–5. Epub 2011/07/09.

18. Tse D, Langston RF, Kakeyama M, Bethus I, Spooner PA, Wood ER, et al. Schemas and memory consolidation. Science. 2007;316(5821):76–82. Epub 2007/04/07.

19. Wang SH, Tse D, Morris RG. Anterior cingulate cortex in schema assimilation and expression. Learn Mem. 2012;19(8):315–8. Epub 2012/07/18.

20. Wang SH, Morris RG. Hippocampal-neocortical interactions in memory formation, consolidation, and reconsolidation. Annu Rev Psychol.2010;61:49–79, C1–4. Epub 2009/07/07.

21. Morris RGM. Elements of a neurobiological theory of hippocampal function: the role of synaptic plasticity, synaptic tagging and schemas. European Journal of Neuroscience. 2006;23(11):2829–46.

22. Carandini M, Churchland AK. Probing perceptual decisions in rodents. Nat Neurosci. 2013;16(7):824–31. Epub 2013/06/27.

23. Whishaw IQ, Tomie J. Of mice and mazes: similarities between mice and rats on dry land but not water mazes. Physiol Behav. 1996;60(5):1191–7. Epub 1996/11/01.

24. Cressant A, Besson M, Suarez S, Cormier A, Granon S. Spatial learning in Long-Evans Hooded rats and C57BL/6J mice: different strategies for different performance. Behav Brain Res. 2007;177(1):22–9. Epub 2006/12/13.

25. Preston AR, Eichenbaum H. Interplay of hippocampus and prefrontal cortex in memory. Current biology: CB. 2013;23(17):R764–73.

26. Haettig J, Sun Y, Wood MA, Xu X. Cell-type specific inactivation of hippocampal CA1 disrupts location-dependent object recognition in the mouse. Learn Mem. 2013;20(3):139–46. Epub 2013/02/19.

27. Assini FL, Duzzioni M, Takahashi RN. Object location memory in mice: pharmacological validation and further evidence of hippocampal CA1 participation. Behav Brain Res. 2009;204(1):206–11. Epub 2009/06/16.

28. Haettig J, Stefanko DP, Multani ML, Figueroa DX, McQuown SC, Wood MA. HDAC inhibition modulates hippocampus-dependent long-term memory for object location in a CBP-dependent manner. Learn Mem. 2011;18(2):71–9. Epub 2011/01/13.

29. Mumby DG, Gaskin S, Glenn MJ, Schramek TE, Lehmann H. Hippocampal damage and exploratory preferences in rats: memory for objects, places, and contexts. Learn Mem. 2002;9(2):49–57. Epub 2002/05/07.

